# Repeatability and Heritability of UAV-Derived Canopy Traits in a Cassava Breeding Population Using Time-Series Data from Two Consecutive Growing Seasons

**DOI:** 10.64898/2026.06.03.729518

**Authors:** Juan Quiros-Vargas, Anna van Doorn, Ismail Y. Rabbi, Hans-Peter Piepho, Juliane Bendig, Wolfgang Zierer, Uwe Sonnewald, Uwe Rascher, Onno Muller

## Abstract

Cassava is a major staple crop in tropical regions, particularly in Sub-Saharan Africa, yet its productivity remains constrained by genetic and agronomic limitations. A major bottleneck in cassava breeding is the difficulty of accurately phenotyping agronomic traits under field conditions using conventional, labor-intensive methods. Here, we evaluated the potential of uncrewed aerial vehicle (UAV)-based phenotyping to quantify canopy growth traits and assess their genetic relevance under realistic field conditions. For this, multi-temporal UAV imagery was collected over two growing seasons (2018-2019 and 2019-2020) in a panel of 46 cassava genotypes planted in fields of the International Institute for Tropical Agriculture (IITA), Nigeria. Canopy height, canopy volume, and their relative growth rates (RGR_h_ and RGR_v_) were extracted at the plot-level, and their seasonal dynamics and canopy-yield relationships were further assessed across developmental stages and environmental conditions. Repeatability (R) and broad-sense heritability (H^2^) were estimated using a linear mixed model (LMM) that partitioned genetic, genotype-by-year, and residual variance components, enabling the evaluation of both measurement reliability and genetic signal. Overall, UAV-derived growth dynamics were found to exhibit comparable patterns across genotypes, reflecting shared seasonal growth trajectories, while canopy-yield relationships varied with developmental stage and environmental conditions. In terms of genetic metrics, R was high for all UAV-derived traits (R = 0.68-0.69), indicating reliable genotype-level assessment across replicates and seasons. In contrast, H^2^ differed substantially among traits. Canopy volume (H^2^ = 0.64) and canopy height (H^2^ = 0.58) exhibited moderate-to-high heritability, reflecting strong genotype effects and comparatively moderate genotype-by-year interactions. However, their relative growth rates showed near-zero H^2^ values, driven primarily by genotype-by-year interaction, indicating a dominant environmental influence. These results demonstrate that UAV-derived canopy height and volume provide a consistent basis for genetic differentiation of cassava genotypes across environments, supporting their use in selection, whereas growth-rate traits are better suited for characterizing growth plasticity and genotype-by-environment interactions.

## 1. Introduction

Cassava is a key staple for food security across tropical regions, especially in Sub-Saharan Africa, where its storage roots provide essential dietary energy (FAO, 2013). Despite its resilience in poor soils and harsh environments, substantial yield gaps persist due to genetic and agronomic limitations (El-Sharkawy, 2004). Breeding varieties with improved productivity is therefore essential to meet rising food demands and strengthen the economic stability of cassava-dependent regions.

One of the primary challenges in cassava breeding is the difficulty of accurately phenotyping agronomic traits in open field conditions. Traditional phenotyping methods for cassava, such as manual height measurements, are time-consuming and may provide less precise and less heritable estimates compared to advanced 3D sensing approaches (Zang et al., 2023). These challenges hinder the efficiency of breeding programs aimed at identifying superior genotypes with enhanced growth and yield potential.

Uncrewed aerial vehicles (UAVs) have emerged as a widely adopted tool in agricultural research, offering a non-destructive, high-throughput approach to plant phenotyping (Bhandari et al., 2023; Guo et al., 2021; Tanaka et al., 2024). UAV-based imaging enables rapid and accurate trait measurements, such as canopy height (Bendig et al., 2015) and biomass accumulation (Quiros Vargas et al., 2019), which are essential for assessing cassava agronomic traits and performance across large breeding trials (Nascimento et al., 2024). Recent work by Gomez Selvaraj et al. (2020) further demonstrated the potential of UAV time series combined with machine learning to predict cassava root yield and extract canopy structure metrics, highlighting UAV phenotyping as a practical selection tool in modern cassava breeding programs. Thus, leveraging UAV-derived data with advanced image processing techniques can facilitate genetic selection and improve breeding efficiency.

Beyond structural canopy traits, UAV time series also enable the extraction of dynamic indicators such as relative growth rates (RGRs), which describe how rapidly plants increase in size relative to their current height. This parameter is widely used in ecology and crop science because it captures temporal patterns of vigor, resource-use efficiency, and developmental pace that static measurements alone cannot reveal (Lambers and Poorter, 1992). When applied in breeding contexts, RGR provides a sensitive measure of how genotypes differ in proportional growth across the season, offering complementary insights into physiological performance and stress responsiveness. Incorporating RGR into UAV-based phenotyping therefore adds a dynamic dimension to the cassava trait assessment, strengthening the ability to characterize genotype performance over time.

The growing availability of UAV-derived data not only overcomes the logistical barriers of traditional cassava phenotyping but also enables a deeper understanding of the genetic and temporal dynamics of plant performance. Moreover, UAV-based time series can help to assess the stability and genetic consistency of key canopy traits, offering valuable insights into their repeatability (R) and heritability (H^2^): two core parameters describing the reliability and genetic basis of the phenotypic variation observed. Repeatability reflects the consistency of trait measurements across environments, replicates, or time points, while heritability quantifies the proportion of phenotypic variance attributable to genetic rather than environmental or residual effects (Schmidt et al., 2019).

High R and H^2^ values of UAV-derived canopy traits indicate that they are both reliable and genetically meaningful, allowing breeders to identify genotypes that combine strong phenotypic performance with stable genetic expression, thereby enhancing selection precision and accelerating cassava improvement. The accurate estimation of these parameters is particularly critical for root crops, where yield and quality traits are strongly influenced by environmental variation and genotype-by-environment interactions. For instance, studies in sweet potato have demonstrated the value of heritability analyses for understanding agronomically important traits and supporting breeding efforts (Gurmu et al., 2018; Mugisa et al., 2022; Sakaigaichi et al., 2022). In cassava, heritability and repeatability analyses have been utilized to understand agronomic traits like root yield and quality, supporting selection decisions (Nascimento et al., 2024; dos Santos et al., 2023). The integration of UAV-based phenotyping with statistical modeling of R and H^2^ therefore represents a powerful framework for accelerating selection, improving genetic gains, and strengthening the efficiency of cassava breeding programs targeting productivity and resilience.

It is essential to consider that while R and H^2^ quantify the genetic contribution to phenotypic variation, their biological interpretation depends on the environmental conditions under which traits are expressed. In cassava, canopy development reflects the coordination between source activity in leaves and sink development in storage roots, processes that are dynamically regulated and highly responsive to climatic variation (Rosado-Souza et al., 2023; Chang and Zhu, 2017). For example, under low water availability, cassava can reduce canopy expansion, potentially altering carbon allocation to storage roots (Muiruri et al., 2021). Consequently, genotype-level differences in canopy traits must be interpreted within intra- and inter-seasonal climatic contexts when evaluating their relevance for root yield formation.

In this study, conducted within the framework of the Cassava Source-Sink (CASS) project, which aims to enhance root yield by increasing assimilate allocation to storage roots (Sonnewald et al., 2020; Zierer et al., 2025), we advance UAV-based phenotyping for cassava by combining high-resolution canopy time series with quantitative genetic analysis to assess the R and H^2^ of key structural UAV-derived canopy traits across multiple seasons. To our knowledge, this study meaningfully advances prior work by providing a multi-season mixed-model variance decomposition of UAV-derived structural canopy traits in cassava, enabling a quantitative assessment of their R and H^2^ across years. Specifically, we aim (i) to quantify the R and H^2^ of canopy height, canopy volume, and their relative growth rates across two consecutive growing seasons in a breeding population of 46 genotypes, and (ii) to evaluate the relationship between high-H^2^ canopy growth traits and root weight across intra- and inter-seasonal climatic contexts, providing insight into environmentally driven modulation of allocation dynamics. With this integrative approach we illustrate the potential of UAV-based time series of canopy traits as a decision-support tool for cassava phenotyping, supporting breeders in the selection of high-performing and genetically stable genotypes.

## 2. Methods

### 2.1. Field experiments

This study was conducted at the International Institute for Tropical Agriculture (IITA), Nigeria, where two field trials were established: one in Ibadan during the 2018-2019 season (Fig. 1a) and another in Ikenne during 2019-2020, about 80 km away in the southwestern region of the country (Fig. 1b). Their location within Nigeria is shown in Fig. 1c. Each trial followed a randomized complete block design with two blocks per season; the block divisions are indicated by dashed white lines in Fig. 1a-b. In each block, a panel of 52 genotypes was initially planted, of which 46 were included in the analysis, as the other 6 genotypes were excluded due to differing randomization procedures. The analyzed genotypes comprised 42 breeding lines and four standard check varieties (TMEB419, IITA-TMS-IBA30572, IITA-TMS-IBA980581, and IITA-TMS-IBA982101). Plots consisted of 20 plants arranged in 5 × 5.5 m rectangles, covering approximately 27.5 m^2^. To minimize border effects, a 1 m inner buffer was applied, resulting in an effective analysis area of 10.5 m^2^.

**Figure 1:**
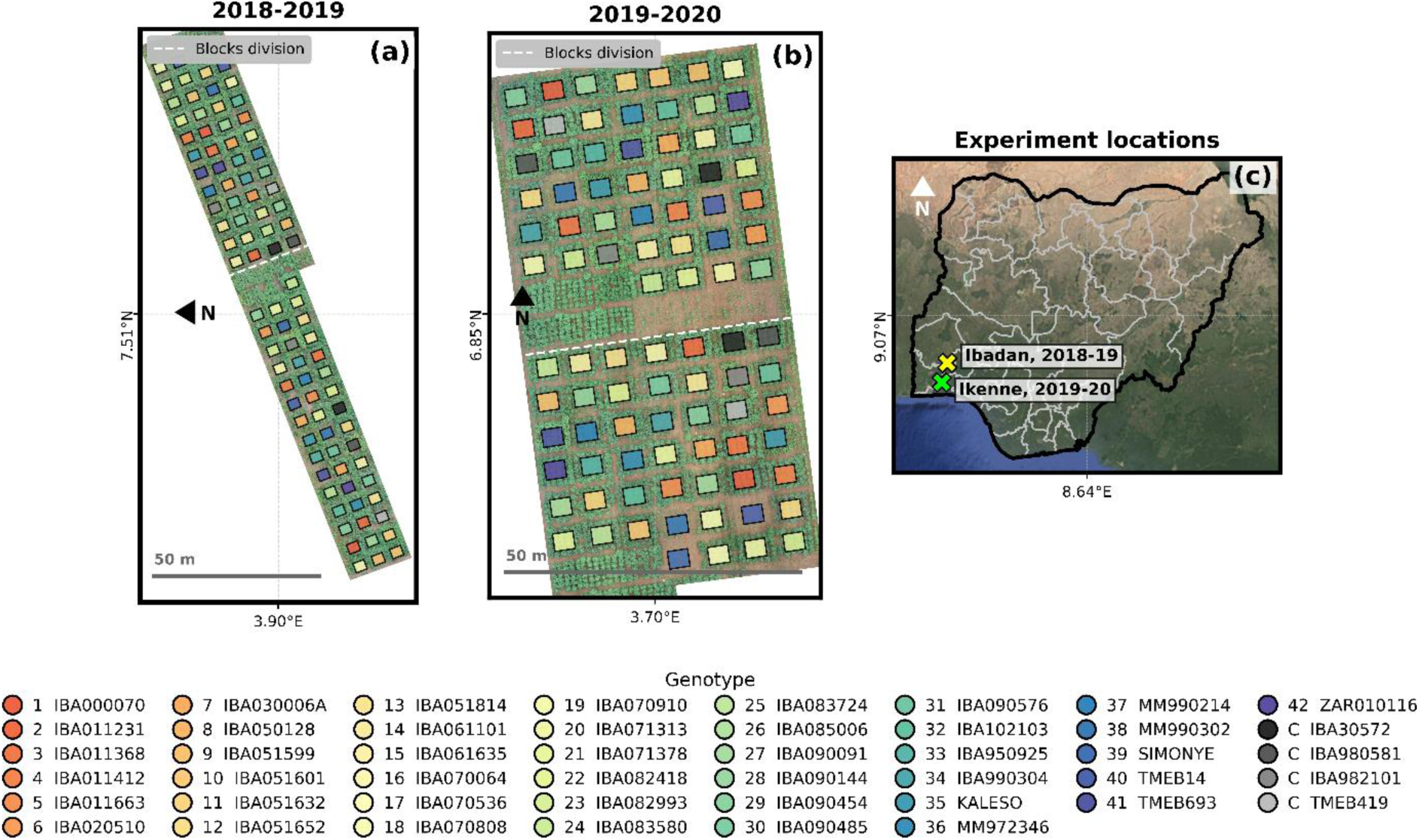
Field experiment locations and plot distributions across two growing seasons. Panels (a) 2018-2019 and (b) 2019-2020 show plot footprints colored by genotype and overlaid on uncrewed aerial vehicle (UAV)-based RGB orthomosaics from 2019-06-17 and 2019-09-06, respectively. Trials were conducted as randomized complete block designs with two blocks per season; block divisions are indicated by dashed white lines. Panel (c) situates the experiments within Nigeria over a Google Satellite Imagery background (google.com/maps) with state and national boundaries. Axes are labeled at the panel centers in geographic coordinates. A 50 m scale bar is shown on the field panels, and the genotype legend appears below. The same color coding and numbering for genotypes are maintained consistently across all figures in this study.

### 2.2. UAV data acquisition

UAV flight campaigns were conducted with a Cinestar-8 (Kopterworx - Kopter d.o.o., Leskovec pri Krškem, Slovenia). A high-resolution RGB camera (Sony alpha 6000 with 35 mm lens; Sony Group Corporation, Tokyo, Japan) was used to collect imagery with 80% overlap (side and forward) at 27 meters above ground level (m AGL), resulting in a pixel size of 0.003 m; however, the images were resampled to 0.010 m for the analysis. Nadir (vertical, top-down views) images were collected close to solar noon, typically between 11:00 and 13:00 h local time at five time points in 2018-2019, and another five time points in 2019-2020 (Table 1). For the March 30, 2020 dataset, approximately 50% of the area was excluded from the final analysis, in order to mitigate the impact of localized blur artifacts within the orthomosaic. To avoid introducing imbalance in the representation of genotypes and plots, this dataset was retained only for descriptive analyses of seasonal dynamics but excluded from the R and H^2^ estimations, which rely on a balanced representation of the experimental design. On average 17 ground control points (GCPs) were used to georeference the data based on their known position measured with a real-time kinematic (RTK)-global navigation satellite system (GNSS) system, achieving an overall accuracy of ~0.03 m.

**Table 1:** Uncrewed aerial vehicle (UAV) data collection dates for the seasons 2018-2019 and 2019-2020.

| *Season* | *Date* | *Season* | *Date* |
| --- | --- | --- | --- |
| 2018-2019 | 19/07/2018 | 2019-2020 | 11/07/2019 |
|  | 17/08/2018 |  | 06/09/2019 |
|  | 17/09/2018 |  | 12/11/2019 |
|  | 14/11/2018 |  | 22/01/2020 |
|  | 17/06/2019 |  | 20/03/2020 |

### 2.3. UAV data processing

Individual raw UAV images were processed with the photogrammetric structure-from-motion software Metashape (Agisoft LLC, St. Petersburg, Russia) to generate georectified orthomosaics and digital elevation models (DEMs) at high spatial resolution (1 cm pixel^−1^), enabling accurate elevation mapping at the individual-plant level (Fig. 2a). The DEMs provide elevations in meters above mean sea level (mAMSL). To obtain crop surface data, the digital terrain model (DTM, also in mAMSL) was subtracted from the DEMs, resulting in crop surface models (CSMs) that represent plant height above ground level (m AGL).

**Figure 2:**
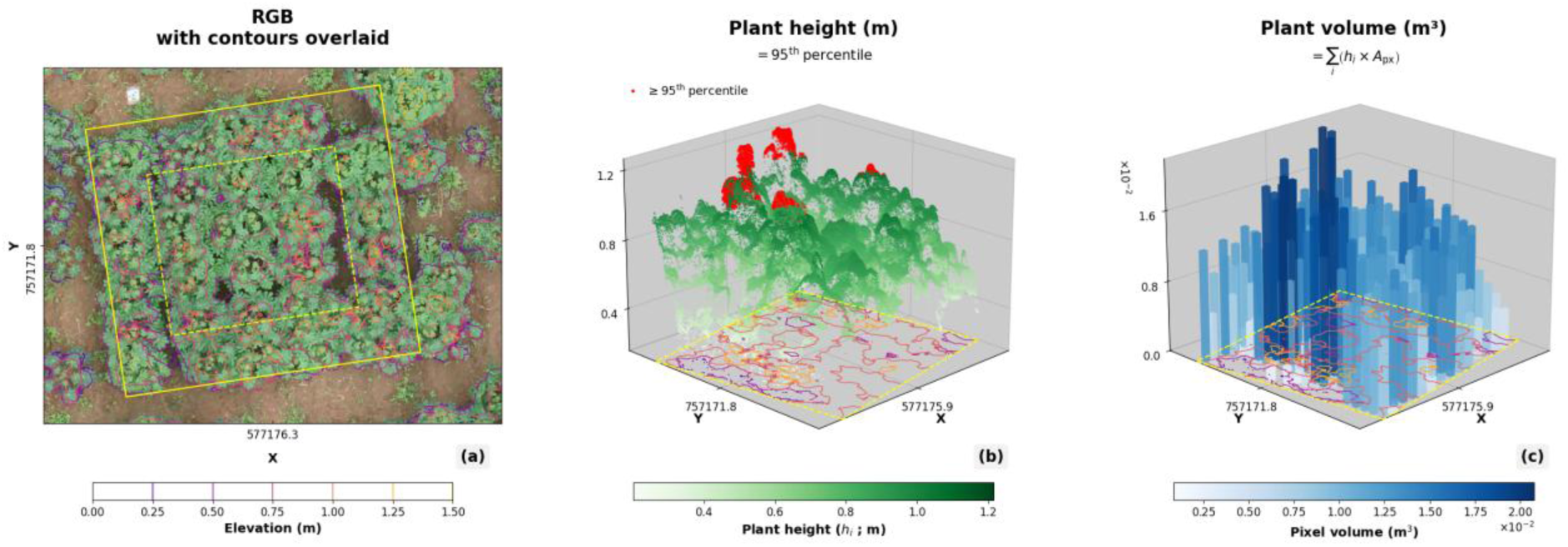
Example of cassava canopy trait extraction from uncrewed aerial vehicle (UAV) data. (a) RGB orthomosaic with elevation contours overlaid. (b) Plant height estimation from the 95th percentile of canopy elevation values; red points indicate pixels at or above this threshold. (c) Plant volume estimation obtained as the sum of pixel-wise volumes (*v_i_* = ℎ*_i_* × *A_px_*), where ℎ*_i_* is canopy height above ground and *A_px_* is the ground area represented by one pixel. The X and Y axes represent UTM coordinates (m), while the Z axes in panels (b) and (c) correspond to plant height (m) and pixel volume (m^3^), respectively. Color scales represent elevation (a), plant height (b), and pixel volume (c).

Plant height was extracted for each plot as the 95^th^ quantile of the CSM values within the 10.5 m^2^ effective analysis area (Fig. 2b), which reduces sensitivity to outliers. Plant volume was computed using eq. 1, as the sum of the CSM pixel-wise heights above ground (*h_i_*), multiplied by the pixel area (*A_px_*; Fig. 2). Ground-truth measurements of plant height were used to validate the UAV-derived estimates. It is important to note that this method of computing canopy volume has limitations, as it becomes less accurate once the cassava canopy closes. From that point in the season onward, it no longer accounts for the side architecture of the plant and instead relies solely on top-of-canopy information, potentially leading to an underestimation of the actual volume. This limitation is further explored in the discussion.

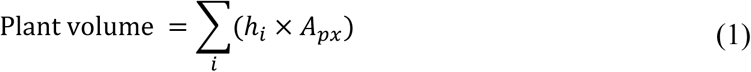

The RGR was calculated as the change in the natural logarithm of plant height or canopy volume between two consecutive time points divided by the corresponding linear time interval (eq. 2). This formulation expresses growth on a proportional basis while preserving time in its original linear scale (Lambers and Poorter, 1992; Lamont et al., 2023). By using logarithmic transformation of the growth variable, the approach reduces the influence of absolute plant size on growth comparisons and enables biologically meaningful evaluation of proportional growth dynamics among genotypes across developmental stages.

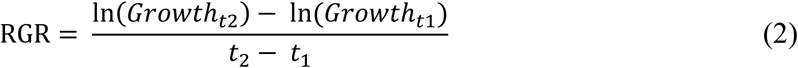

### 2.4. Seasonal dynamics

Time series of the growth traits (height, volume, RGR_h_ and RGR_v_) were plotted alongside regional precipitation and temperature data acquired from the NASA POWER Project website (https://power.larc.nasa.gov/). The raw weather information was downloaded as a grid of sampling points, each spaced 56 km apart, as illustrated in Annex 1. For 2018-2019 the weather information was averaged across the three closest points to Ibadan (Annex 1b), whereas for 2019-2020 it was across the three closest points to Ikenne Annex 1c.

In addition to full-season trajectory plots, canopy-yield relationships were evaluated at three representative UAV acquisition dates per season (early, mid, and late or mid-to-late). For each target date, canopy trait values were first averaged across plot-level measurements to obtain a single value per genotype. Genotypes were then ranked according to the respective trait and partitioned into ten consecutive groups of 4 to 5 genotypes sharing similar trait magnitude. Group-level medians of canopy traits and root weight were subsequently used to evaluate canopy-yield relationships through Pearson correlation analysis.

### 2.5. Estimation of repeatability (R) and heritability (H^2^)

Field trials are subject to environmental heterogeneity, incomplete replication, and spatial variation; therefore, heritability should be interpreted as a measure of precision in distinguishing genetic from environmental sources of variation under realistic field conditions (Schmidt et al., 2019). Following this framework, we estimated R and H^2^ of UAV-derived canopy traits using a linear mixed model (LMM) that partitions variance components attributable to genotype, genotype-by-year interaction, and residual effects. Trait values were analyzed on the plot-mean scale: for each genotype i, year *j*, and replicate plot *k*, UAV-derived measurements across all days after planting (DAPs) were averaged to obtain a single response per plot and year, thereby avoiding explicit modeling of serial correlation over time. These plot-level means (*y_ijk_*) were then analyzed using an LMM with a fixed effect for year and random effects for genotype, genotype-by-year interaction, and block (year × replicate):

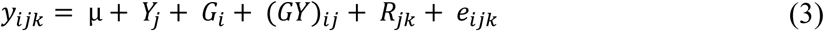

where *y_ijk_* is the response of the *i^th^* genotype in the *k^th^* replicate within the *j^th^* year; µ is the overall mean; *Y_j_* is the fixed main effect of year; 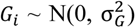 is the random genetic effect with variance 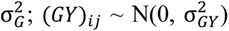 is the random genotype-by-year interaction with variance 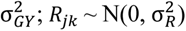 is the random block effect for the *j^th^* year and *k^th^* replicate with variance 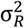; and 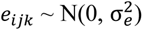 is the residual at the plot-mean level with variance 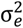. Variance components were estimated by restricted maximum likelihood (REML). The *R_jk_* was included in the model to account for design-related and micro-environmental variation among plots. Based on the fitted variance components, R at the plot-level was defined as the ratio of explained plot-to-plot variance (genetic + genotype-by-year) to total plot-to-plot variance:

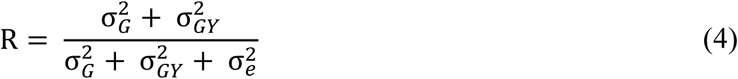

Broad-sense heritability on genotype means across years and replicates was computed by scaling the interaction and residual variances by the numbers of years and plots contributing to the genotype mean. Let *n_y_* denote the number of years and *n_r_* the effective number of replicates per genotype. Then

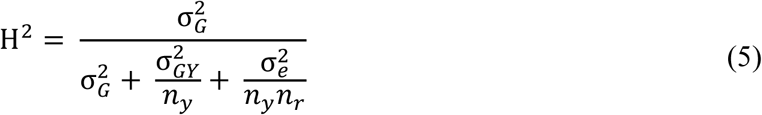

The term 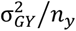 reflects averaging genotype performance across years, whereas 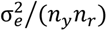 reflects averaging across the total number of plots contributing to genotype means over years and replicates. Residual variation 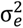 captures within-genotype plot variation not explained by the random terms, including measurement error and micro-environmental heterogeneity at the plot-mean level. The variance component of *R_jk_* was not included in R and H^2^ estimations because it represents design-specific, non-genetic variation among replicate plots within years.

High R and H^2^ values indicate that UAV-derived canopy traits are both reliable and genetically meaningful, allowing breeders to identify genotypes that combine strong phenotypic performance with stable genetic expression, thereby enhancing selection precision and accelerating cassava improvement.

Figure 3 presents the overall workflow from UAV data acquisition to the computation of R and H^2^.

**Figure 3:**
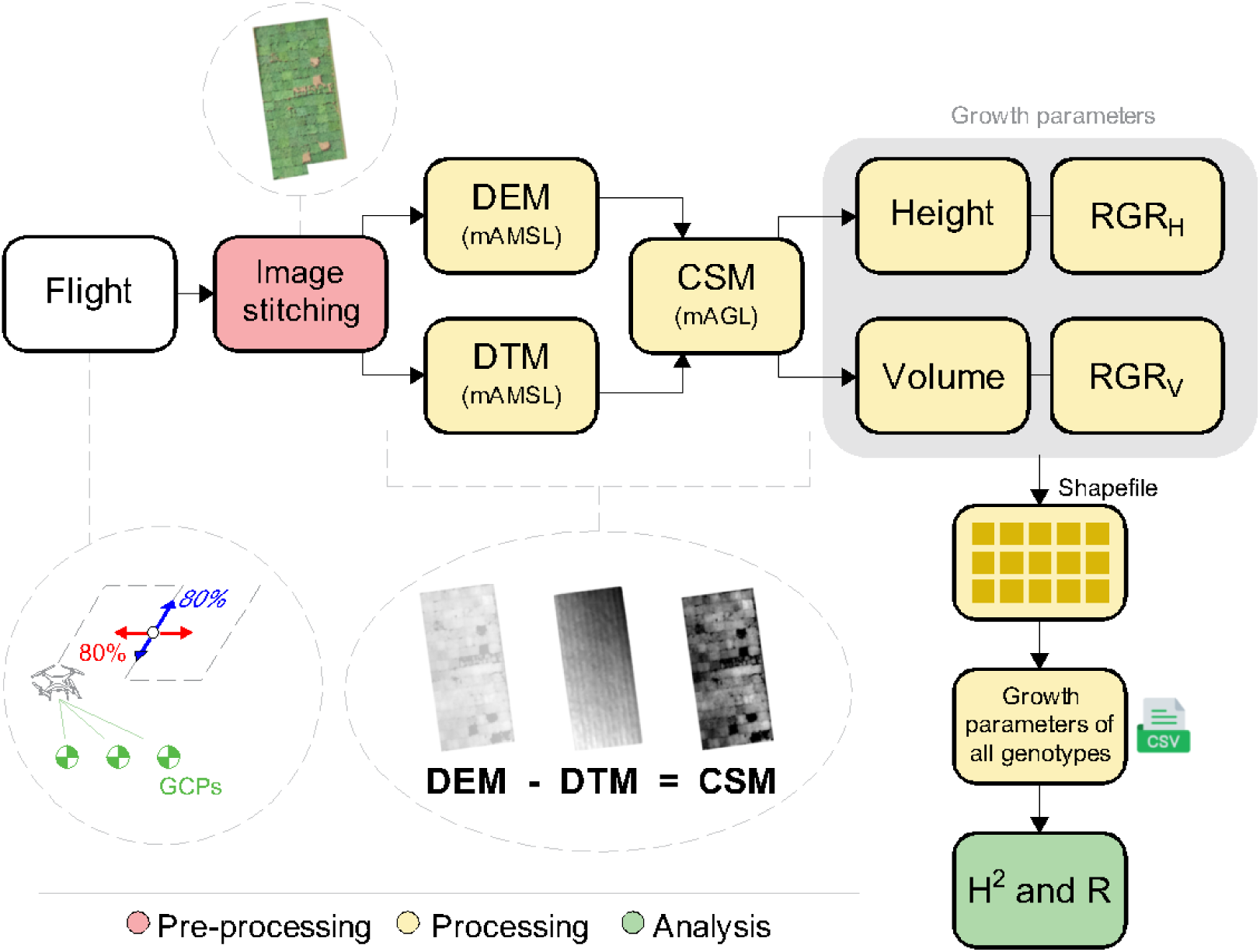
Workflow for extracting growth parameters from uncrewed aerial vehicle (UAV)-based imagery. After the flight, images are stitched to generate a digital elevation model (DEM) and a digital terrain model (DTM). The crop surface model (CSM) is derived as DEM - DTM, representing vegetation height. Growth parameters (height, volume and their relative growth rates (RGR_h_ and RGR_v_) are extracted and linked to genotype data using a shapefile, then saved as a CSV file for calculating heritability (H^2^) and repeatability (R). Color codes represent pre-processing (red), processing (yellow), and analysis (green) steps.

### 2.6. Bootstrap-based estimation of uncertainty in R and H^2^

For each trait, uncertainty in R and H^2^ was quantified using a design-consistent parametric bootstrap procedure. For this, the fitted mixed model was used to generate synthetic datasets by simulating new random effects for Genotype, Genotype×Year, and Year×Rep (block), plus residual errors, while preserving the original Genotype×Year×Rep design. In this procedure, each bootstrap replicate retained the same experimental structure (same genotypes, years, and number of replicates and plots per combination) but replaced the observed values with new, randomly simulated phenotypes that followed the same statistical model and variance structure estimated from the real data. Each simulated dataset was refitted using the same model, and new estimates of R and H^2^ were obtained. The SE of these estimates across 100 bootstrap replicates was used to quantify the uncertainty of R and H^2^.

### 2.7. Genotype-level canopy-yield association analysis

To evaluate how UAV-derived canopy traits relate to root yield performance at the genotype-level, we quantified genotype-wise mean canopy height, canopy volume, and root weight for both growing seasons (2018-2019 and 2019-2020). The four standard check varieties (TMEB419, IBA30572, IBA980581, and IBA982101) were grouped into a single control class, represented by the average of their canopy and yield values within each season.

To assess the joint variation between canopy traits and yield across genotypes, we generated two-dimensional scatterplots (UAV-derived canopy trait versus root weight) for each season separately and partitioned each plot into four quadrants defined by the median canopy trait and median root weight. This approach allowed classification of phenotypes into four groups: (I) small shoot, high yield; (II) large shoot, high yield; (III) large shoot, low yield; and (IV) small shoot, low yield.

Additionally, kernel density estimation (KDE) contours were computed using a Gaussian kernel to visualize the multivariate density of genotype-level points and highlight global trends in the canopy-yield relationships. This analysis provides a framework for identifying architectural types with distinct biomass allocation patterns and for evaluating the consistency of these relationships across contrasting seasonal conditions.

### 2.8. Assessment of genomic relatedness among genotypes

In order to provide a genetic context for the observed phenotypic variation in UAV-derived canopy traits, genomic relatedness among genotypes was assessed using a genomic relationship heatmap. For this, a genome-wide single nucleotide polymorphism (SNP) data were filtered for minor allele frequency (MAF > 5%) and used to compute a pairwise genetic distance matrix. In addition, a hierarchical clustering was applied to this matrix to generate dendrograms, which were used to structure the heatmap and visualize pairwise genetic similarity among genotypes.

## 3. Results

### 3.1. Validation of UAV-Derived Plant Height

Ground-truth data from 14 genotypes were available to validate the accuracy of UAV-derived plant growth measurements. For this, we assessed the correlation between UAV data and field measurements across the two consecutive growing seasons. Fig. 4 shows the relationships between UAV-derived plant height and manual measurements of shoot weight and plant height for the 2018-2019 and 2019-2020 seasons, respectively. Shoot weight (as a proxy of plant size) was used as the ground-truth trait for the 2018-2019 in the absence of manual plant height measurements for that year.

**Figure 4:**
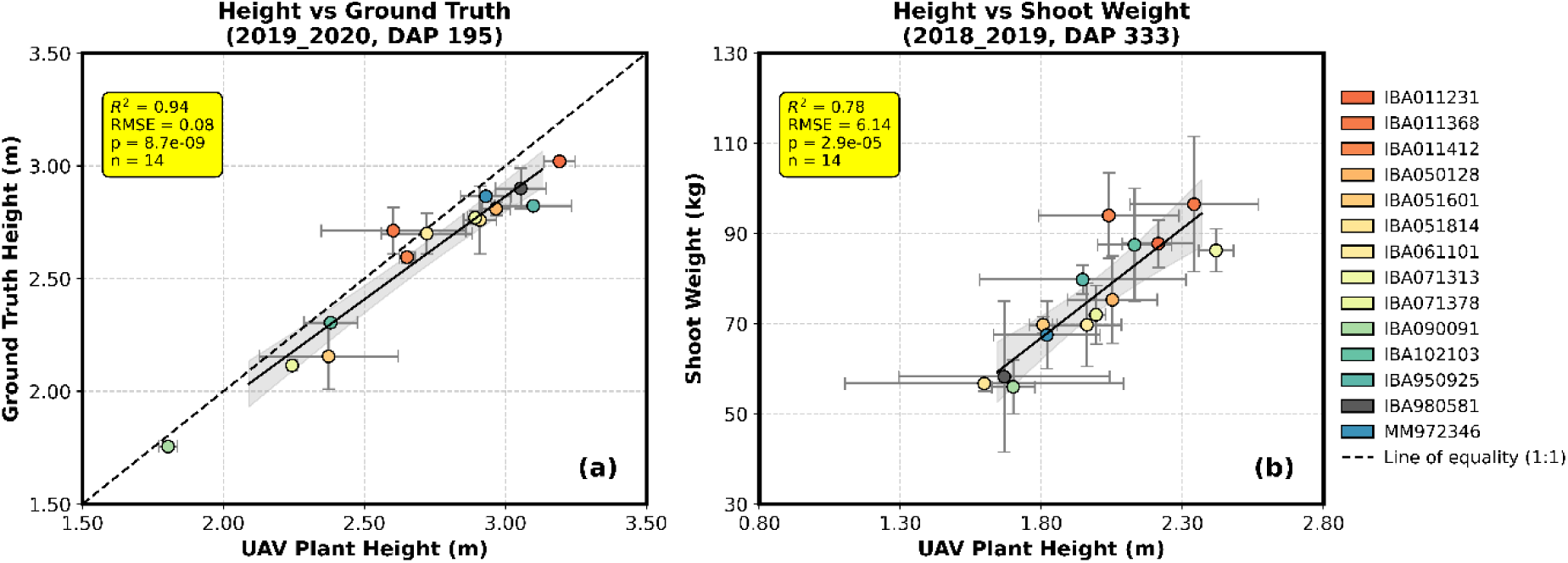
Relationships between uncrewed aerial vehicle (UAV)-derived plant height and ground-truth measurements across two consecutive growing seasons. Panel (a) shows the correlation between UAV-derived plant height and manually measured ground-truth plant height during the 2019-2020 season (DAP 195, closest date to the manual height measurements). Panel (b) presents the correlation between UAV-derived plant height and shoot weight measured at harvest during the 2018-2019 season (DAP 333, closest available date to harvest), used as a proxy due to the absence of manual plant height data for that season. Yellow boxes summarize the statistical performance of each linear regression, including the coefficient of determination (R^2^), root mean square error (RMSE), p-value, and sample size (n, as the number of plot means used in the regression), reflecting the strength, accuracy, and significance of each relationship. Error bars represent the standard error of the mean for each trait within a plot (calculated from replicate plants), while shaded areas around the regression lines indicate the 95% confidence intervals of the predicted values, providing a visual representation of model uncertainty. The dashed line in panel (a) represents the line of equality (1:1), included as a reference for perfect agreement between UAV-based and ground-truth height measurements. Genotype color coding corresponds to that introduced in Figure 1.

A strong correlation of R^2^ = 0.89 (RMSE = 0.16 m, p < 0.01) between UAV-derived plant height and manually measured plant height for 2019-2020 (Fig. 4a) validates the accuracy of UAV-derived height estimation. Similarly, the strong association between UAV-derived plant height and shoot weight for 2018-2019 (R^2^ = 0.78, RMSE = 74.64 g, p < 0.01; Fig. 4b) indicates that UAV-derived canopy height was closely related to aboveground biomass in that season.

### 3.2. Seasonal Dynamics of UAV-Derived Plant Height and Volume

Across both growing seasons, UAV-derived plant height and canopy volume exhibited comparable seasonal trajectories (Fig. 5a-d). In 2018-2019, both height (Fig. 5a) and volume (Fig. 5b) increased steadily during the first four months of the season. Toward the end of the season, both traits approached a plateau, with slight declines observed in a subset of genotypes, indicating a deceleration in canopy expansion that was particularly evident as a reduction in canopy volume in some genotypes. In 2019-2020, plant height (Fig. 5c) increased overall in a sustained manner, with a brief mid-season slowdown before reaching a peak in the mid- to late season, after which it stabilized toward the final measurement dates. In contrast, canopy volume (Fig. 5d) increased substantially during the early and mid-season but declined markedly during the final two months, suggesting a decoupling between vertical growth and canopy structural development at this stage.

**Figure 5:**
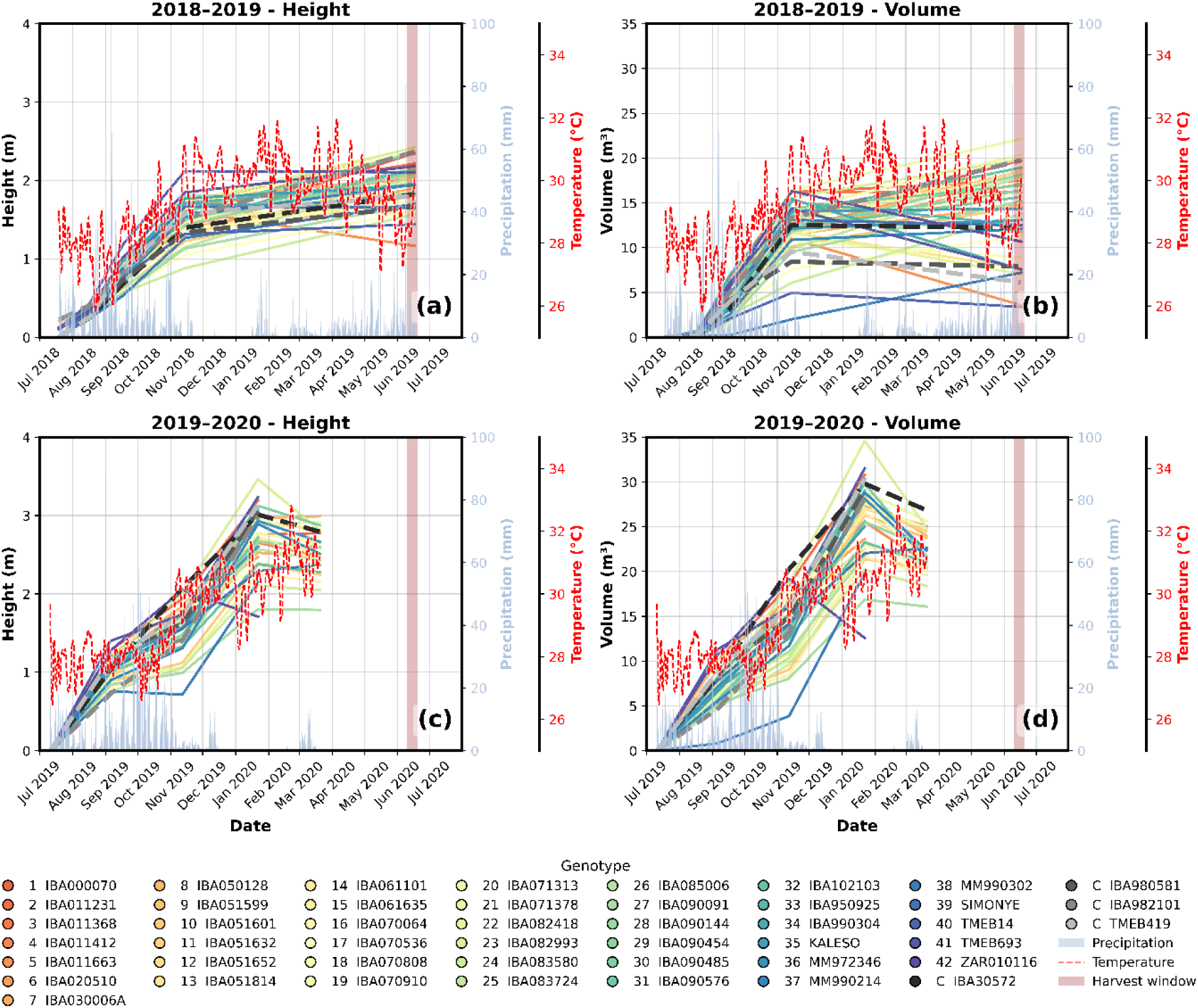
Seasonal dynamics of uncrewed aerial vehicle (UAV)-derived plant height and canopy volume across two growing seasons (2018-2019 and 2019-2020). Panel (a) shows height trends for 2018-2019, panel (b) shows volume trends for 2018-2019, panel (c) shows height trends for 2019-2020, and panel (d) shows volume trends for 2019-2020. Colored solid lines represent trajectories of the 46 individual genotypes (see legend). Precipitation is displayed as light blue bars (right y-axis), and temperature as a red dashed line representing a 7-day moving average (secondary right y-axis). The shaded vertical band indicates the approximate harvest window. The x-axis represents calendar date. Genotype color coding and numbering correspond to those introduced in Fig. 1.

Weather information provided valuable context for interpreting these growth patterns. During the early to mid-season months (July to November), frequent rainfall events combined with gradually rising temperatures created favorable conditions for vigorous canopy development across genotypes in both seasons. In 2018-2019, as rainfall declined and temperatures stabilized or slightly decreased toward the end of the season, growth slowed and both plant height and canopy volume approached a plateau, with volume trajectories diverging more strongly among genotypes. In contrast, the late phase of the 2019-2020 season was characterized by notably reduced precipitation combined with elevated temperatures. Under these conditions, plant height largely stabilized, whereas canopy volume declined, suggesting a reduction in canopy density under increasingly dry and warm late-season conditions. Notably, relative to the standard check varieties in both height and volume, a larger proportion of genotypes outperformed the controls in 2018-2019, whereas under the warmer and drier conditions of 2019-2020, most genotypes showed reduced performance compared to the controls.

These patterns indicate that UAV-derived traits consistently capture seasonal growth dynamics across years while revealing trait-specific responses at distinct developmental stages and under varying climatic conditions, thereby providing a sensitive framework for investigating genotype-by-environment interactions in field settings.

### 3.3. Seasonal Dynamics Of UAV-Derived Relative Growth Rates of Height and Volume

Figure 6 illustrates the seasonal dynamics of UAV-derived relative growth rates for plant height (RGR_h_) and canopy volume (RGR_v_) across the 2018-2019 and 2019-2020 growing seasons, together with precipitation and temperature trends. In 2018-2019, both RGR_h_ (Fig. 6a) and RGR_v_ (Fig. 6b) exhibited a pronounced early-season peak during August to October, reflecting rapid proportional canopy expansion shortly after establishment. Following this initial surge, both RGR metrics declined progressively throughout the season, indicating a gradual reduction in proportional canopy expansion over time. While absolute height and volume continued to increase during mid-season, their relative growth rates progressively decreased.

**Figure 6:**
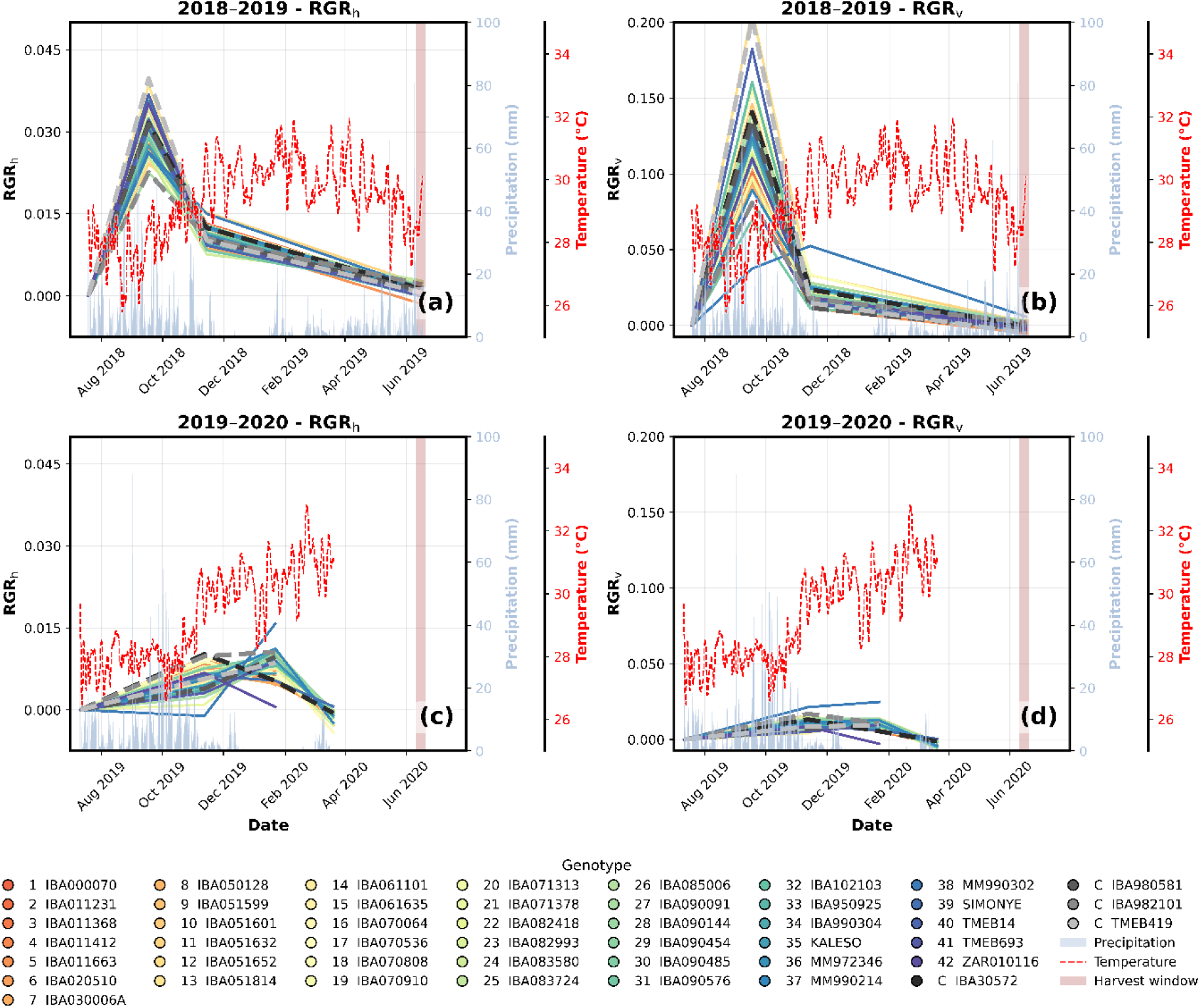
Seasonal dynamics of uncrewed aerial vehicle (UAV)-derived relative growth rate for height (RGR_h_) and volume (RGR_v_) across two growing seasons (2018-2019 and 2019-2020). Panel (a) shows RGR_h_ trends for 2018-2019, panel (b) shows RGR_v_ trends for 2018-2019, panel (c) shows RGR_h_ trends for 2019-2020, and panel (d) shows RGR_v_ trends for 2019-2020. Colored solid lines represent trajectories of the 46 individual genotypes (see legend). Precipitation is displayed as light blue bars (right y-axis), and temperature as a red dashed line representing a 7-day moving average (secondary right y-axis). The shaded vertical band indicates the approximate harvest window. The x-axis represents calendar date. Genotype color coding and numbering correspond to those introduced in Fig. 1.

The season of 2019-2020 showed a distinctly different temporal pattern (Fig. 6c-d). Early-season RGR_h_ and RGR_v_ were comparatively modest, with most genotypes exhibiting gradual proportional increases during July-October 2019, consistent with slower initial canopy development relative to the previous season. A moderate mid-season increase then emerged between approximately November 2019 and January-February 2020, particularly evident for RGR_h_ (Fig. 6c), where several genotypes exhibited increased relative height growth during mid-season. Following this mid-season peak, both RGR_h_ and RGR_v_ declined toward near-zero values late in the season. These patterns demonstrate that relative growth metrics captured inter-annual differences in growth timing and intensity, revealing shifts from predominantly early-season growth in 2018-2019 to relatively more delayed mid-season growth dynamics in 2019-2020. In contrast to the patterns observed for absolute canopy traits, the relative growth rates of the standard check varieties did not exhibit a consistent tendency to outperform or underperform the broader genotype panel in either season, but rather fell within the general range of variation observed across genotypes.

Notably, across both seasons, a single genotype (MM990302) exhibited distinct RGR seasonal dynamics relative to all other genotypes, characterized by peaks at different time points for both height and volume, while maintaining comparatively low to intermediate canopy size.

### 3.4. Temporal Dynamics of the Relationship Between Plant Height and Storage Root Weight

Figure 7 shows the relationship between UAV-derived plant height and storage root weight at different developmental stages across the two growing cycles, together with the corresponding precipitation and temperature profiles. Clear differences in the temporal dynamics of this relationship emerged between years.

**Figure 7:**
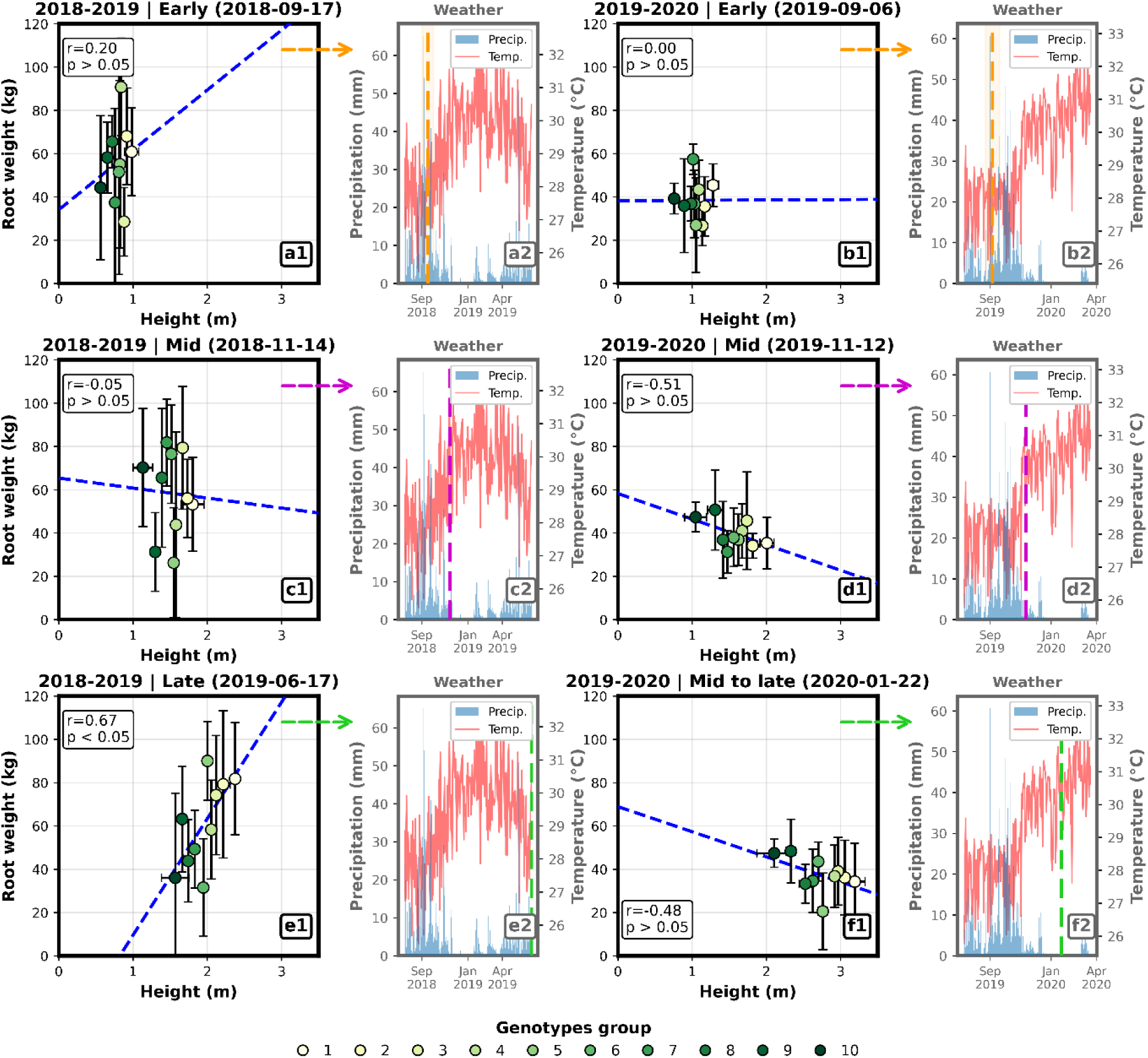
Phase-specific canopy-yield relationships contextualized by seasonal weather across two contrasting growing seasons. Scatterplots (a1-f1) illustrate the relationship between storage root weight and the uncrewed aerial vehicle (UAV)-derived canopy height at three representative phenological phases per season (2018-2019: Early, Mid, Late; 2019-2020: Early, Mid, and Mid-to-late). For each target date, plot-level canopy height was averaged per genotype to obtain a single value, genotypes were ranked by canopy height and partitioned into ten consecutive groups of similar magnitude, and group-level medians of canopy height and root weight were used to calculate the Pearson’s correlation coefficient (r) and associated p-values. Points represent group medians, and error bars denote within-group standard deviation. Dashed blue lines indicate linear regression fits. Weather panels (a2-f2) display daily precipitation (blue bars) and 7-day moving-average air temperature (red line) over the corresponding season. Colored dashed vertical lines mark the UAV acquisition date associated with each correlation panel (arrows indicate panel pairing). A corresponding figure showing the same phase-specific analysis using canopy volume instead of height is provided in Annex 2.

During the 2018-2019 cycle, the early-stage assessment (September 2018) revealed only a weak and non-significant association between plant height and storage root weight (r = 0.20, p > 0.05). At this stage, genotypic differences in canopy height were not yet reflected in differences in root biomass. A similarly weak and non-significant relationship was observed at mid-season (November 2018; r = −0.05, p > 0.05), indicating that although canopy development progressed, variation in aboveground growth still did not systematically translate into storage root accumulation. By contrast, at the late-season time point (June 2019), the relationship shifted markedly. A significant positive correlation emerged between plant height and root weight (r = 0.67, p < 0.05), with taller genotypes consistently exhibiting higher storage root biomass.

A different pattern was observed in 2019-2020. At the early stage (September 2019), plant height showed no association with storage root weight (r = 0.00, p > 0.05). At mid-season (November 2019), a moderate, although non-significant, negative relationship emerged (r = −0.51, p > 0.05). This negative tendency persisted into the mid-to-late stage (January 2020; r = −0.48, p > 0.05), suggesting that, from mid-season onward, taller genotypes tended to exhibit lower storage root weight.

These results demonstrate an inter-annual shift in biomass allocation dynamics. Whereas one cycle showed a progressive alignment between canopy height and storage root accumulation toward the end of the season, the other exhibited a negative association during key developmental stages.

### 3.5. Repeatability (R) and Heritability (H^2^)

Table 2 summarizes the variance partitioning and corresponding estimates of R and H^2^ for the four UAV-derived canopy traits. For canopy height, the genotypic variance component 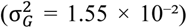 represents the largest share (~44%) of the total phenotypic variance, indicating a strong and consistent genetic signal across seasons. The genotype-by-year interaction component 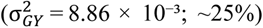 contributes a smaller but still relevant fraction, reflecting moderate season-dependent modulation of height expression. Residual variance 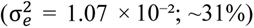 remains relatively low. A similar pattern is observed for canopy volume, where 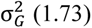 again constitutes the largest component (~43%), in this case followed by 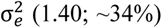 and then 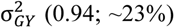 The substantial genetic contribution indicates clear genotypic differentiation in canopy volume, while genotype-by-year interaction and residual variance play secondary roles.

**Table 2.** Variance components estimated for uncrewed aerial vehicle (UAV)-derived traits across two seasons. Values correspond to the genetic variance 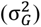, permanent environmental variance attributable to genotype-by-year 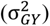, and residual variance 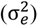 obtained from linear mixed models fitted to per-plot raw trait means for plant height, canopy volume, RGR_h_, and RGR_v_. Percentages in parentheses indicate the relative contribution of each variance component to the total phenotypic variance within the respective trait (summing to 100%).

| *Trait* | $\sigma_G^2$ | $\sigma_{GY}^2$ | $\sigma_e^2$ |
| --- | --- | --- | --- |
| Height | $1.55 \times 10^{-2}$ (44.1%) | $8.86 \times 10^{-3}$ (25.3%) | $1.07 \times 10^{-2}$ (30.6%) |
| Volume | $1.73 \times 10^0$ (42.6%) | $9.37 \times 10^{-1}$ (23.0%) | $1.40 \times 10^0$ (34.4%) |
| RGR<sub>h</sub> | $5.17 \times 10^{-15}$ (~0%) | $9.86 \times 10^{-7}$ (63.6%) | $5.63 \times 10^{-7}$ (36.4%) |
| RGR<sub>v</sub> | $1.16 \times 10^{-14}$ (~0%) | $7.01 \times 10^{-5}$ (68.3%) | $3.25 \times 10^{-5}$ (31.7%) |

In contrast, the relative growth-rate traits exhibit markedly different variance structures. For RGR_h_, the genotype-by-year interaction component 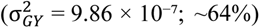 clearly exceeded the residual variance 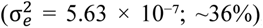, while the genotypic variance 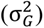 was essentially zero, indicating that proportional height growth was almost exclusively influenced by seasonal and environmental conditions, with no detectable genetic component. A similar pattern was observed for RGR_v_, where 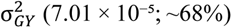 represented the largest share of total variance, followed by 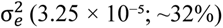, and with genotypic variance again near zero. These results demonstrate that relative growth dynamics for both traits are almost entirely driven by genotype-by-environment modulation and residual variation, with no meaningful genetic signal. Consequently, despite moderate R, H^2^ for RGR_h_ and RGR_v_ is effectively zero, reflecting their extreme environmental sensitivity and lack of cross-seasonal genetic stability, in contrast to the absolute canopy traits.

The variance component patterns in Table 2 demonstrate that UAV-derived canopy height and volume capture strong, stable genetic signals with limited residual noise, whereas relative growth-rate traits primarily reflect environmentally driven dynamics. These contrasting variance structures mechanistically explain the observed differences in H^2^ among traits and underscore the complementary roles of static structural traits and dynamic growth metrics in UAV-based cassava phenotyping.

Figure 8 presents repeatability (R; Fig. 8a) and heritability (H^2^; Fig. 8b) estimates for UAV-derived canopy height, volume, RGR_h_, and RGR_v_, with bootstrap-derived standard errors reflecting the uncertainty of these estimates. Repeatability values were consistently moderate to high across all traits, ranging from 0.66 to 0.69, indicating relatively stable trait expression across replicates and measurement periods within seasons. In contrast, H^2^ values differed substantially among traits, reflecting marked differences in the relative contribution of genetic and environmental sources of variation. Canopy volume showed the highest H^2^ (0.64), followed by canopy height (H^2^ = 0.58), consistent with the variance structures described in Table 2, where genotypic effects accounted for a substantial proportion of total phenotypic variation and genotype-by-year interactions remained comparatively moderate.

**Figure 8:**
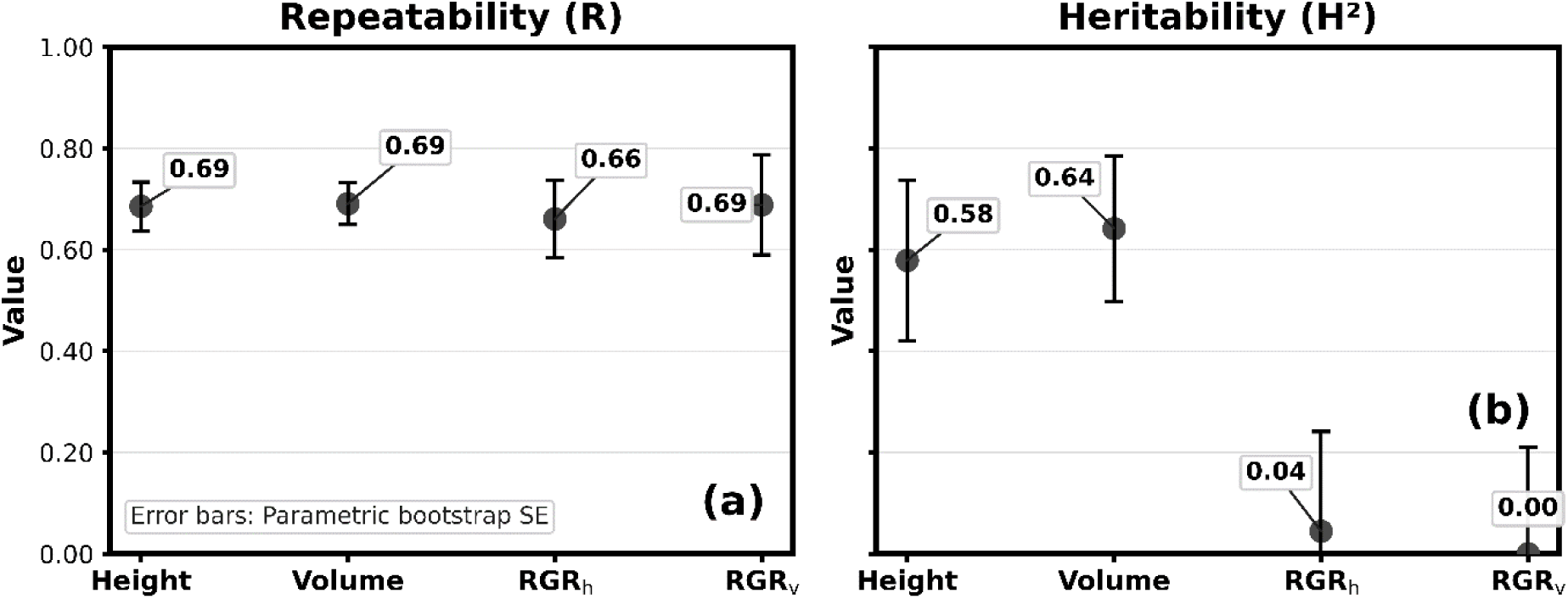
(a) Repeatability (R) and (b) heritability (H^2^) for uncrewed aerial vehicle (UAV)-derived canopy traits: Height, Volume, RGR_h_ (relative growth rate of height), and RGR_v_ (relative growth rate of volume). Error bars represent bootstrap standard errors (SE) obtained from 100 design-consistent parametric bootstrap replicates. For each trait, synthetic datasets were generated by simulating new random effects and residuals under the fitted mixed model (preserving the original Genotype-Year-Rep design) and refitted to estimate the variability of R and H^2^. The resulting SEs reflect uncertainty consistent with the experimental design and the hierarchical sampling of phenotypic variation.

By comparison, the relative growth-rate traits exhibited very low H^2^ values. RGR_h_ approached zero heritability (H^2^ = 0.04), whereas RGR_v_ exhibited effectively zero heritability (H^2^ ≈ 0.00), consistent with their variance partitioning being dominated by genotype-by-year interaction and residual components rather than stable genotypic effects. Despite these low H^2^ values, repeatability for both RGR traits remained moderate (R = 0.66-0.69), indicating that relative growth dynamics retained some degree of within-season consistency despite strong environmental modulation across years. Bootstrap-derived standard errors were generally smaller for R than for H^2^, and within H^2^, estimates were more precise for height and volume than for the two RGR traits. These results highlight the suitability of static UAV-derived canopy traits for genetic evaluation, while indicating that proportional growth-rate traits capture highly environment-dependent aspects of plant performance with limited cross-seasonal genetic stability.

### 3.6. Genotypic Variation in Shoot-Root Growth Balance Across Seasons

Building on the moderate-to-high H^2^ observed for canopy height and volume, Figure 9 examines how genotype-level canopy size relates to storage root yield at representative late-season time points in two contrasting growing cycles. A quadrant-based framework is used to highlight distinct allocation strategies, separating genotypes characterized by coordinated shoot-root growth from those exhibiting contrasting source-sink dynamics.

**Figure 9:**
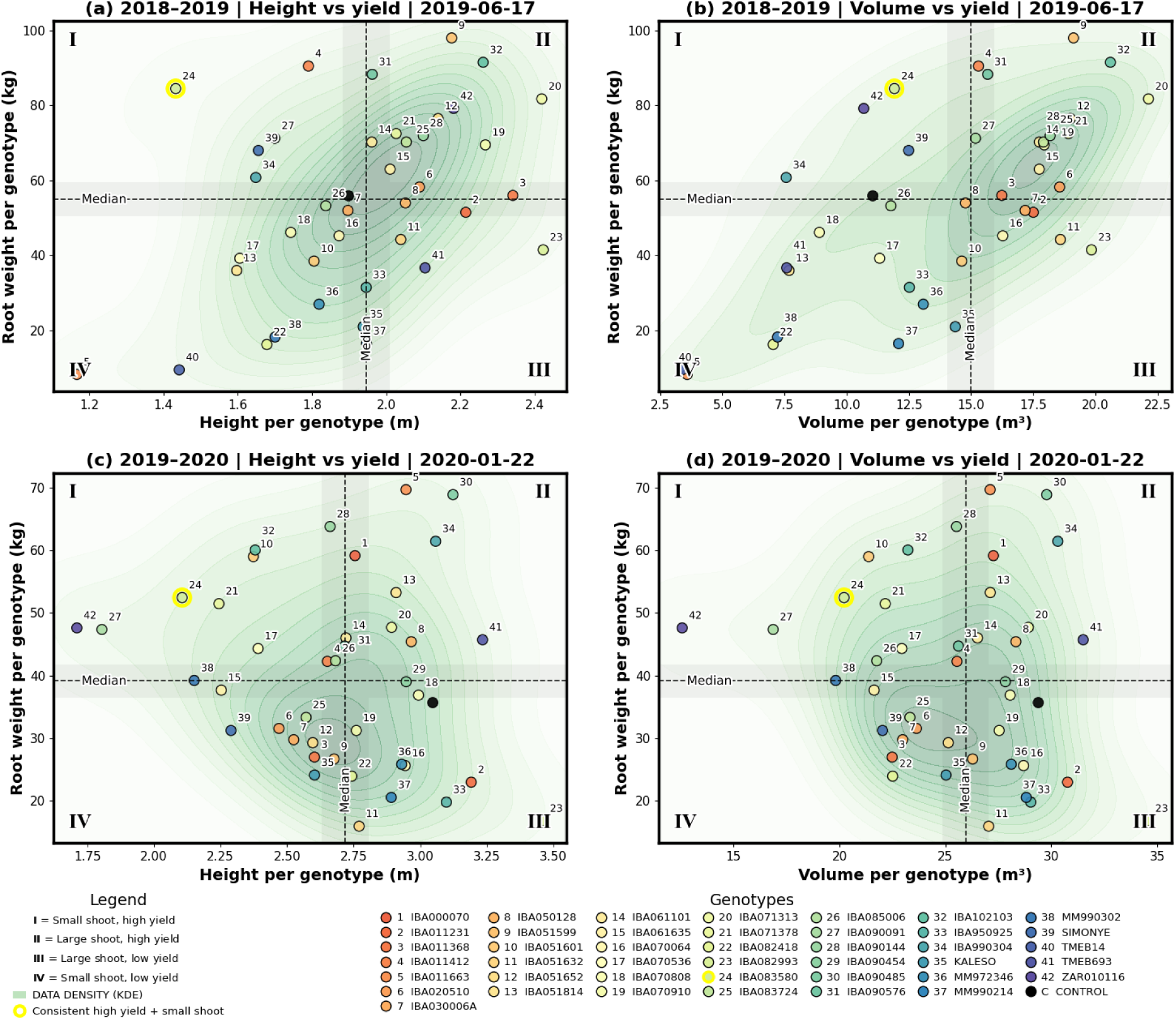
Genotype-level relationships between canopy size and storage root yield at representative late-season time points in two contrasting growing seasons. Scatterplots show genotype-mean canopy height (a, c) and canopy volume (b, d) versus genotype-mean storage root weight for 2018-2019 (2019-06-17) and 2019-2020 (2020-01-22). Each point represents the mean per genotype at the specified date. Background shading indicates data density (kernel density estimation, KDE). Dashed vertical and horizontal lines denote the medians of canopy size and root weight, defining four quadrants: I, small shoot-high yield; II, large shoot-high yield; III, large shoot-low yield; and IV, small shoot-low yield. The yellow-highlighted point indicates the genotype that consistently combined relatively small canopy size with high yield across seasons. Point colors and numeric identifiers correspond to those introduced in Fig. 1.

In 2018-2019 (panels a and b), both canopy height and canopy volume show a generally positive association with storage root yield. Most genotypes cluster along an upward density gradient, as emphasized by the KDE background, indicating that larger canopy size is typically associated with higher root biomass in this season. The median-based quadrant partitioning reveals that approximately two thirds of the high-yielding genotypes fall within Quadrant II (large shoot-high yield), reflecting coordinated shoot and root development. In contrast, only approximately 10-15% of genotypes appeared in Quadrant I (small shoot-high yield), indicating relatively efficient storage root production despite more limited canopy size.

In contrast, the 2019-2020 panels (c and d) display a markedly different structure. The distribution of points is more dispersed and lacks the clear positive alignment observed in the previous season. High-yielding genotypes are distributed across both small- and large-canopy quadrants, and the density gradient is more centralized rather than strongly diagonal. This pattern reflects the weaker or altered relationship between canopy size and yield under the environmental conditions prevailing in this cycle.

A particularly illustrative case is the genotype IBA083580 (highlighted with a yellow halo), which was consistently located in Quadrant I (small shoot-high yield) across all panels, suggesting a phenotype potentially associated with a more efficient source-sink balance.

## 4. Discussion

Across both growing seasons, UAV-derived growth dynamics followed broadly similar temporal trajectories across genotypes, with the exception of MM990302, which showed a distinct pattern in relative growth rates. This shared behavior is in line with the genomic relationship heatmap (Annex 3), which indicates a relatively shallow genetic structure within the breeding population. In this context, the moderate level of genetic divergence may partially explain the convergence in seasonal shoot development dynamics, as closely related genotypes are expected to exhibit comparable structural traits. These findings are consistent with previous work demonstrating that UAV time-series phenotyping can capture genetically controlled variation in plant growth traits (Ye et al., 2023), with our results further showing that this signal can be detected even within relatively narrow phenotypic ranges across a genetically cohesive population.

The contrasting behavior observed between absolute growth values and their RGRs when comparing the standard check varieties with the broader genotype panel highlights that these metrics capture fundamentally different dimensions of plant performance. While canopy height and volume represent cumulative structural outcomes that integrate growth over time, relative growth rates describe changes in growth that vary according to specific environmental conditions (Lambers and Poorter, 1992). As a consequence, differences in final canopy size between controls and the general population are not necessarily driven by transient differences in growth rate, but rather by how biomass is allocated and maintained across the season. Thus, the final plant size emerges as the result of both the duration of growth and the partitioning of assimilates among plant organs (Poorter et al., 2012). This distinction emphasizes that cumulative growth outcomes provide a more informative basis for genotype comparison under field conditions.

The variation in the relationship between canopy height and storage root yield across developmental stages and environmental conditions, as shown in Fig. 7, suggests that canopy size is a context-dependent predictor of yield, reflecting dynamic adjustments in shoot-root biomass allocation. This interpretation is consistent with Gomez Selvaraj et al. (2020), who showed that the strength of canopy-based predictions of cassava root biomass depends on phenological stage, with later developmental phases providing the most reliable associations. From a physiological perspective, these patterns align with the framework described by El-Sharkawy (2004), in which cassava yield is governed by the balance between source activity and sink strength. The author further describes that, under favorable conditions, larger canopies can sustain higher assimilate supply and support storage root bulking, whereas under stress, increased canopy size may impose greater transpirational demand and maintenance costs, potentially limiting carbon allocation to storage roots. This supports the view that canopy-yield relationships in cassava are inherently dynamic and must be interpreted within both temporal and environmental contexts.

These growth dynamics formed the basis for the estimation of repeatability and heritability in our study and, in contrast to previous contributions (e.g., Nascimento et al., 2024), enable, to the best of our knowledge, the first multi-season genetic evaluation of UAV-derived canopy traits in cassava under field conditions. Within this framework, we found that a substantial proportion of the phenotypic variance (approximately 65%) was attributable to genotype-dependent effects (i.e., genotype and genotype-by-year interaction), which is reflected in consistently high repeatability estimates. In this regard, it is worth noting that the concept of repeatability has been less explored in UAV-based studies targeting structural traits; for example, Smith et al. (2024) report repeatability as a within-season metric describing the consistency of biomass predictions across growth stages and experiments. In contrast, our multi-season framework estimates R in a genetic context, capturing the stability of genotype performance across years.

For H^2^, the genotype-by-year interaction component is excluded from the numerator of the equation, in contrast to R, in order to isolate the purely genetic contribution to phenotypic variation. Under this framework, canopy height and volume still exhibit moderate-to-high H^2^ values, indicating that these traits can be estimated with greater genetic precision, highlighting their potential as robust targets for selection in UAV-based cassava phenotyping based on structural traits. In contrast, the corresponding growth-rate traits (RGR_h_ and RGR_v_) show substantially lower (near 0) H^2^, which indicates that a larger share of their variance is driven by genotype-by-environment interactions. With this, our results demonstrate the value of multi-season UAV phenotyping for disentangling stable genetic effects from environment-dependent variation, positioning structural canopy traits as reliable indicators for genetic evaluation while underscoring that growth-rate metrics capture complementary, environmentally responsive dimensions of plant performance.

Although moderate to high, the H^2^ values observed in our study are lower than those reported for remotely sensed plant height in other crops such as wheat, where heritability has approached around 0.90 under both conventional open-field conditions in variety trials (Zang et al., 2023) as well as under more controlled conditions using a structured field phenotyping platform (Madec et al., 2017). In cassava, comparable H^2^ values have been reported for UAV-derived vegetation indices by Nascimento et al. (2024), who observed moderate-to-high heritability (0.51-0.94) using RGB and multispectral imagery. However, their single-season analysis likely contributed to these relatively high estimates by limiting environmental variability and genotype-by-environment interactions. In contrast, our multi-season approach provides a more comprehensive assessment of UAV-derived heritability by explicitly capturing temporal environmental variation and its influence on trait expression, thereby offering a more robust evaluation of trait stability under field conditions.

While previous studies have predominantly focused on vegetation indices or conventional field measurements, our integration of UAV-based canopy time series with mixed-model variance partitioning across two growing seasons extends this framework to structural canopy traits. When compared with multi-season high-throughput field phenotyping of physiological traits in durum wheat reported by dos Santos et al. (2021), whose H^2^ values for chlorophyll fluorescence traits ranged from approximately 0.40 to 0.74 depending on year, trait, and drought conditions, our estimates for canopy height (0.58) and canopy volume (0.64) fall within a comparable range. This alignment suggests that UAV-derived structural traits can provide a level of genetic information comparable to more specialized physiological measurements, supporting their relevance as reliable targets for selection.

The moderate-to-high heritability observed for canopy height and volume further emphasizes the relevance of absolute growth traits for genotype selection, particularly within breeding frameworks such as the CASS consortium, where improvement targets include enhanced carbohydrate partitioning toward storage roots (Sonnewald et al., 2020). In this sense, Figure 9 indicates that the relationship between canopy development and storage root yield is not fixed, but varies across environmental conditions, highlighting the dynamic nature of source-sink interactions. Notably, the genotype IBA083580, which repeatedly exhibited a small shoot size combined with relatively high yield, suggests that certain biomass allocation patterns may have a genetic component. However, other genotypes closely related to IBA083580, according to the genomic relatedness analysis (Annex 3), did not exhibit the same canopy-yield relationship. This observation reinforces that, although genetic factors contribute to source-sink dynamics, these relationships remain strongly influenced by environmental conditions, reflecting the interaction between genetic factors and environmental plasticity (Rosado-Souza et al., 2023).

While the present study demonstrates the potential of UAV-derived structural traits for assessing genetic variability in cassava, several aspects could be improved to increase the precision of R and H^2^ estimates of growth traits computed from aerial imagery. As emphasized by Schmidt et al. (2019), the accuracy of these parameters is highly dependent on the experimental design, particularly the number of replications and environments evaluated, since greater replication reduces error variance and enhances the reliability of genotype mean separation. Moreover, temporal consistency in repeated measurements is equally relevant (Tanaka et al., 2024), as irregular intervals compromise reliable stage-specific estimation of R and H^2^, thereby shifting the analytical focus toward season-integrated trait values. Such time-integrated estimates are prone to inflating residual variance unrelated to genetic effects and limit the detection of stage-specific genetic differentiation. Therefore, future studies should aim to increase the number of replications and maintain homogeneous flight intervals to minimize residual variability and obtain more robust and comparable estimates of R and H^2^ across environments.

Despite the strong performance of UAV-derived height and volume traits for cassava breeding, a remaining limitation concerns the incomplete representation of canopy volume when the canopy closes. This occurs at later growth stages, when overlapping plant crowns obscure the sub-canopy region, causing part of the plant structure below the upper canopy surface to be missed in nadir-derived 3D reconstructions (Annex 4). Quirós Vargas et al. (2025; poster at EPPS 2025) propose to address this issue by integrating complementary side-view imagery to correct the top canopy information with sub-canopy structural details. This limitation highlights an important methodological frontier that warrants development in a dedicated study aimed at improving individual-plant volume reconstruction and its application in cassava phenotyping.

## 5. Conclusions

This study demonstrates the value of multi-temporal UAV-based phenotyping for the genetic evaluation of structural canopy traits in cassava under field conditions. Across two contrasting growing seasons characterized by differences in temperature and precipitation, canopy height and canopy volume showed consistently high R and moderate-to-high H^2^, indicating that these traits provide reliable and genetically informative indicators for genotype differentiation across environments. In contrast, relative growth rate traits exhibited lower H^2^, reflecting a stronger influence of genotype-by-environment interactions and highlighting their relevance for capturing growth plasticity rather than stable genetic effects.

The integration of temporal canopy dynamics revealed that canopy-yield relationships are not static, but vary with developmental stage and environmental conditions. From a physiological perspective, this context dependency reflects the dynamic balance between source activity and sink strength in cassava and highlights the importance of considering both phenological timing and environmental variability when interpreting UAV-derived traits for selection.

Overall, our findings establish UAV-derived canopy height and volume as robust traits for genetic evaluation in cassava breeding, while highlighting the complementary role of growth-rate metrics in characterizing genotype responsiveness to environmental conditions. The multi-season framework presented here provides a more realistic assessment of trait stability and genetic signal under field conditions than single-environment studies, and represents a step forward toward integrating UAV-based phenotyping into operational cassava breeding pipelines.

Future work could build on these findings by improving the temporal consistency of UAV acquisitions and increasing the number of replications, as well as by advancing 3D canopy reconstruction methods to more comprehensively capture canopy structure, particularly after canopy closure. These improvements would further enhance the accuracy and applicability of UAV-derived traits for high-throughput phenotyping and selection in cassava.

## Acknowledgements

This work was funded through grants to Prof. Uwe Sonnewald by the Bill & Melinda Gates Foundation (Grant number 008053) and Gates Agricultural Innovations (Grant number 58147). The conclusions and opinions expressed in this work are those of the author(s) alone and shall not be attributed to the Foundation. Under the grant conditions of the Foundation, a Creative Commons Attribution 4.0 License has already been assigned to the Author Accepted Manuscript version that might arise from this submission. Please note works submitted as a preprint have not undergone a peer review process.

Open access is funded by the Deutsche Forschungsgemeinschaft (DFG, German Research Foundation) - 491111487.

## Annexes

**Annex 1:**
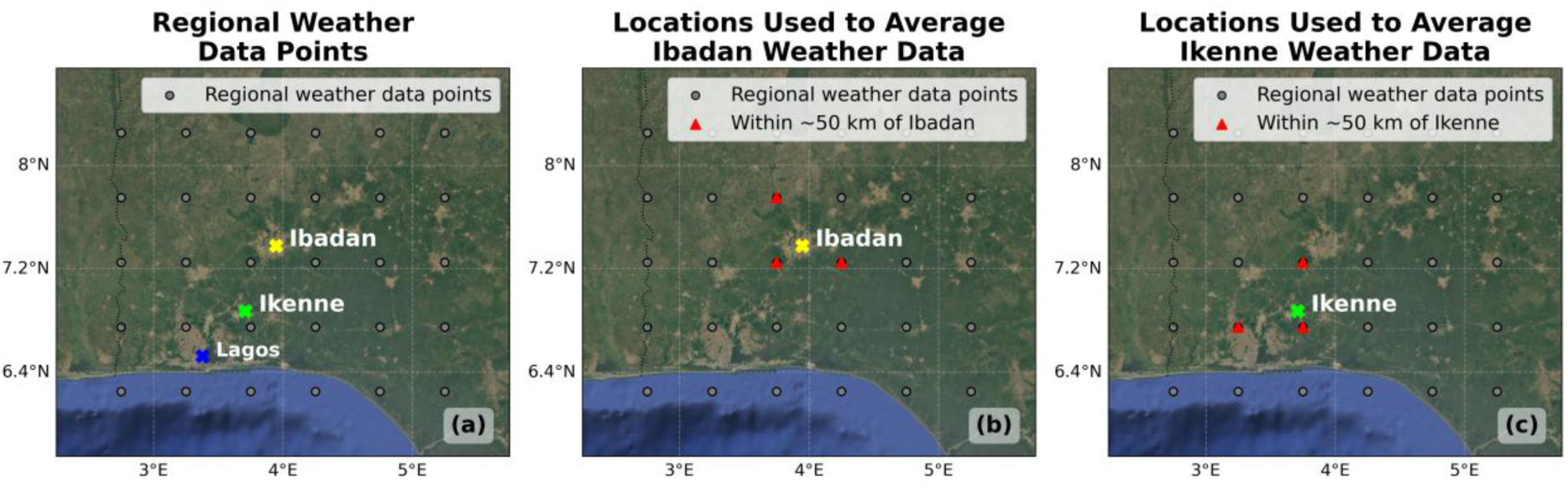
Regional weather data points downloaded from the NASA POWER Project website (https://power.larc.nasa.gov/; black circles) across the study area, with Ibadan (yellow star), Ikenne (green square; a); Lagos (blue star) is presented for geographical reference, as the biggest city around the experiments. Weather stations within ~50 km of Ibadan (b) and Ikenne (c; red triangles) used to calculate the averaged information for each location. The maps are projected in the geographic coordinate system (latitude and longitude, WGS 84). The background satellite imagery is sourced from Google Satellite Imagery (google.com/maps).

**Annex 2:**
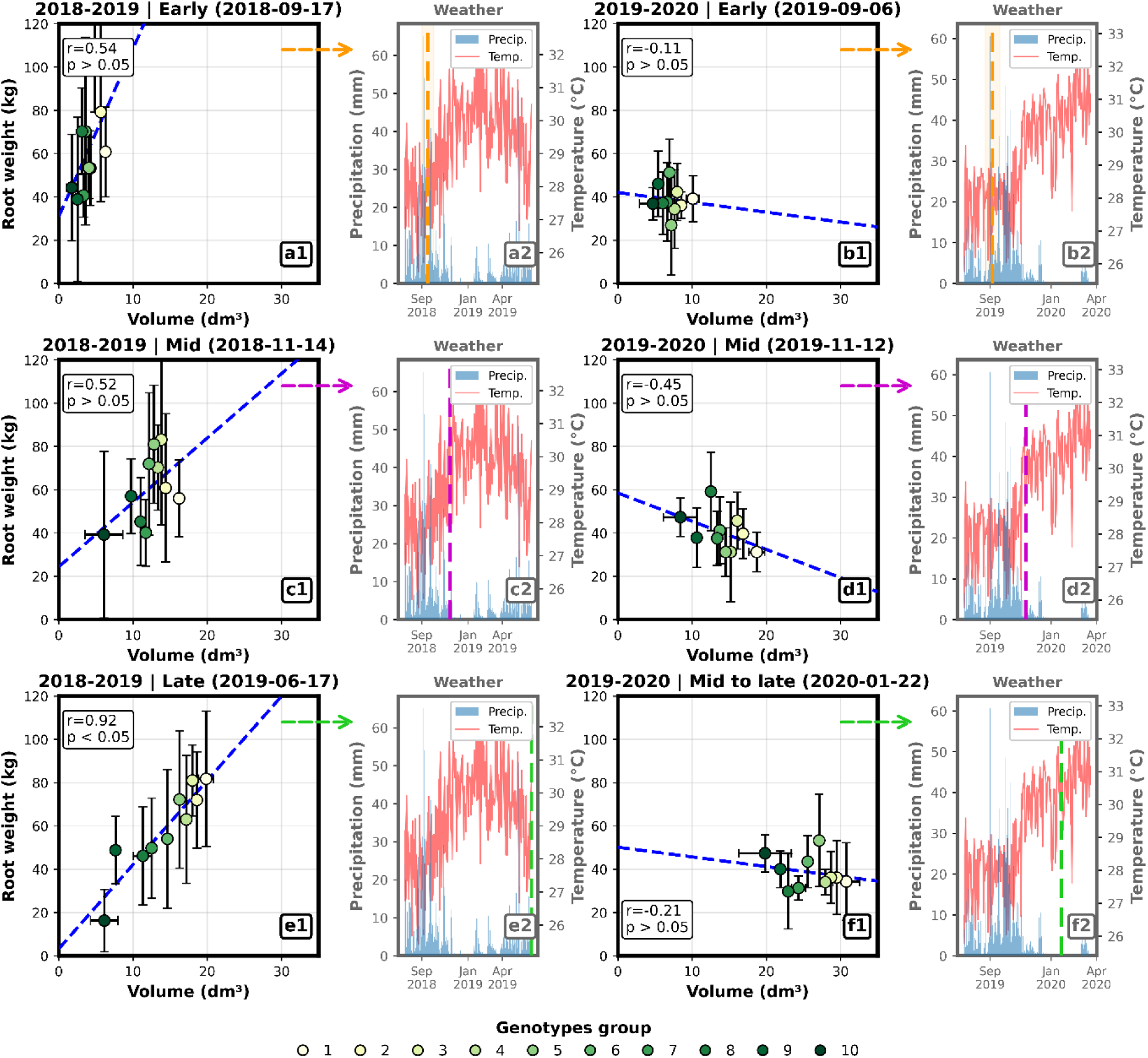
Phase-specific canopy-yield relationships contextualized by seasonal weather across two contrasting growing seasons. Scatterplots (a1-f1) illustrate the relationship between storage root weight and uncrewed aerial vehicle (UAV)-derived canopy volume at three representative phenological phases per season (2018-2019: Early, Mid, Late; 2019-2020: Early, Mid, and Mid-to-late). For each target date, plot-level canopy volume was averaged per genotype to obtain a single value, genotypes were ranked by canopy volume and partitioned into ten consecutive groups of similar magnitude, and group-level medians of canopy volume and root weight were used to calculate the Pearson’s correlation coefficient (r) and associated p-values. Points represent group medians, and error bars denote within-group standard deviation. Dashed blue lines indicate linear regression fits. Weather panels (a2-f2) display daily precipitation (blue bars) and 7-day moving-average air temperature (red line) over the corresponding season. Colored dashed vertical lines mark the UAV acquisition date associated with each correlation panel (arrows indicate panel pairing).

**Annex 3:**
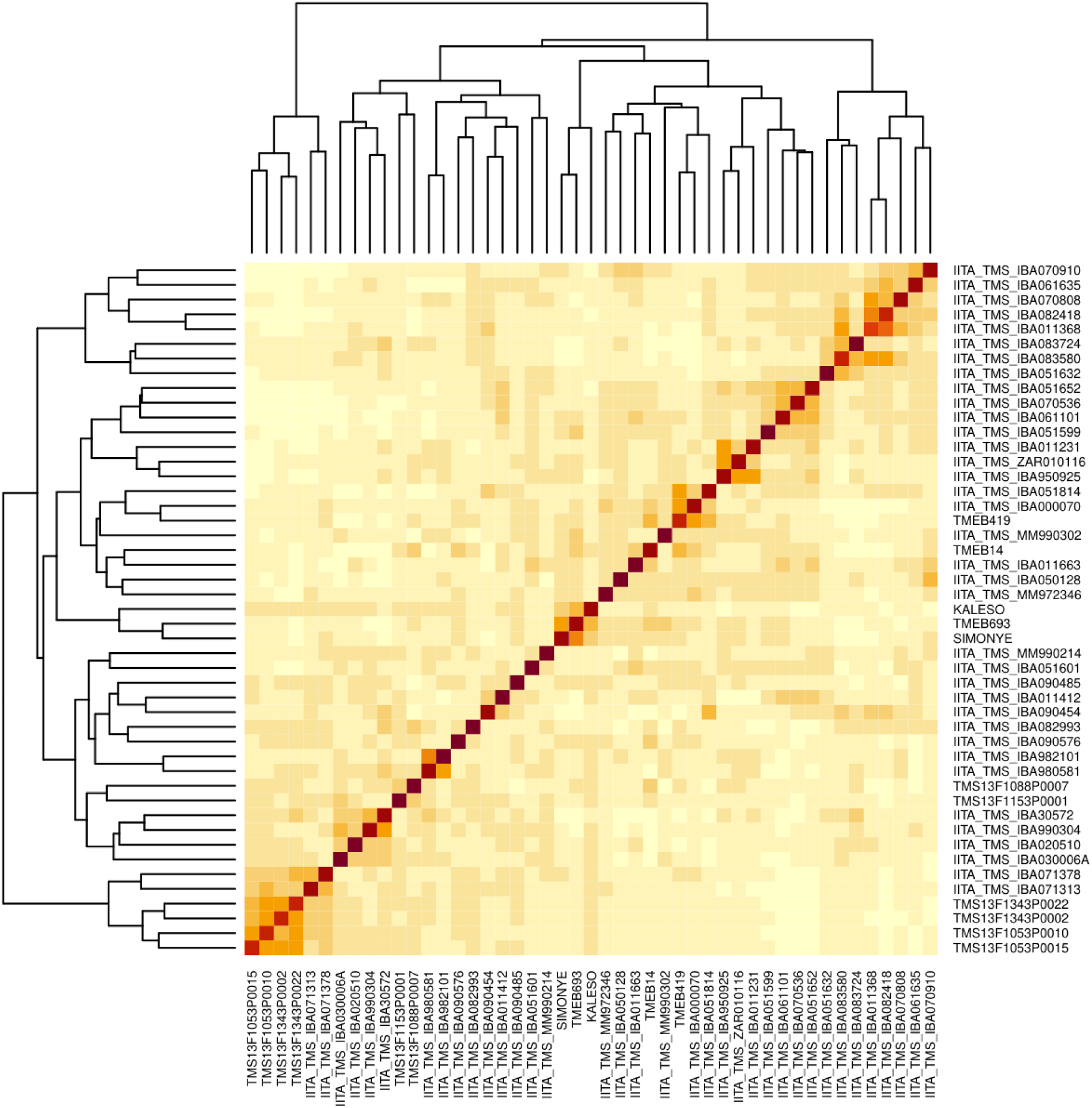
Genomic relationship heatmap showing pairwise genetic similarity among cassava genotypes based on genome-wide single nucleotide polymorphism (SNP) data filtered by minor allele frequency (MAF > 5%); the color scale represents the degree of similarity between genotype pairs, with warmer colors (yellow to red) indicating higher similarity and lighter tones indicating lower similarity. The diagonal corresponds to self-comparisons (maximum similarity), while off-diagonal patterns reflect relative genetic distances among genotypes. Dendrograms derived from the same distance matrix illustrate the clustering structure of the population. The heatmap comprises the full set of 52 genotypes originally planted in the field experiment, including the six genotypes excluded from the main analysis due to differing randomization procedures, corresponding to those starting with “TMS13F1”.

**Annex 4:**
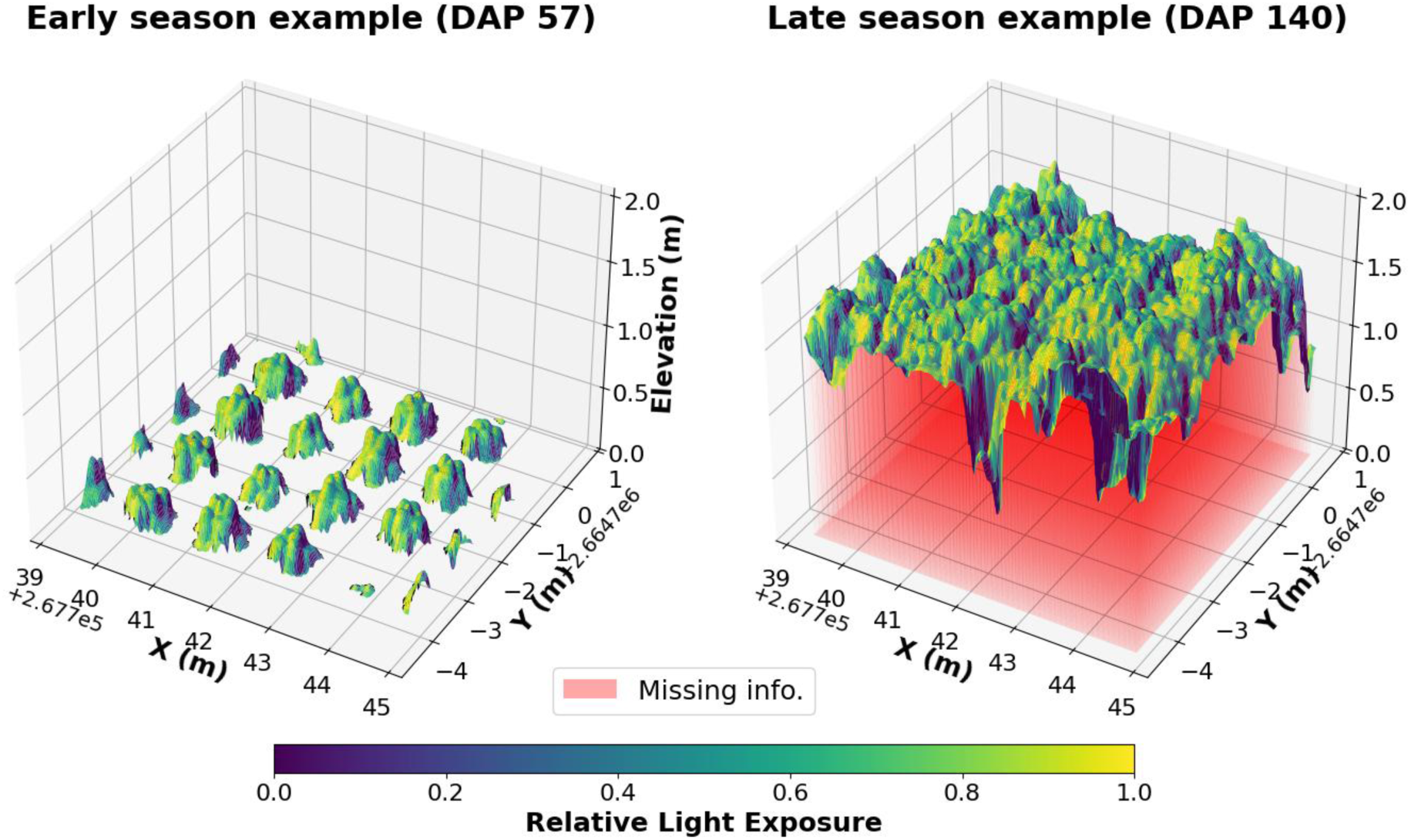
Early- and late-season examples of cassava canopy elevation models derived from UAV-based crop surface models (CSMs). The left panel illustrates the early growth stage (DAP 57), while the right panel shows the late stage (DAP 140), where overlapping canopies result in areas of missing information below the upper canopy surface (highlighted in red). Color gradients represent relative light exposure derived from canopy surface orientation. Adapted from Quiros-Vargas et al. (2025).

